# Generation of a new Tbx6-inducible reporter mouse line to trace presomitic mesoderm derivatives throughout development and in adults

**DOI:** 10.1101/2020.12.10.419275

**Authors:** Laurent Yvernogeau, Anna Klaus, Carina van Rooijen, Catherine Robin

## Abstract

The presomitic mesoderm (PSM) is initially an unsegmented structure localized on each side of the neural tube of the developing embryo, which progressively segments to form the somites. The somites will segregate and partition to generate the dorsal dermomyotome and the ventral sclerotome. Endothelial and myogenic cells of both the trunk and limbs are derived from the somites. There is a lack of efficient reporter mouse models to label and trace the PSM derivatives, despite their crucial contribution to many developmental processes. In this study, we generated a tamoxifen inducible transgenic Tbx6 mouse line, Tg(Tbx6_Cre/ERT2)/ROSA-eYFP, to tag and follow PSM-derivatives from early embryonic stages until adulthood. After induction, endothelial and myogenic cells can be easily identified within the trunk and limbs with proper expression patterns. Since our Tg(Tbx6_Cre/ERT2)/ROSA-eYFP model allows to permanently label the PSM-derived cells, their progeny can be studied at long-term, opening the possibility to perform lineage tracing of stem cells upon aging.

## INTRODUCTION

During embryonic development, endothelial cells (ECs) and myogenic cells (MCs) of the body (epaxial domain) and limbs (hypaxial domain) are derived from the somites. Somites are segments located along the antero-posterior axis of the embryo that are formed after segregation of the presomitic mesoderm (PSM). The somite ultimately forms two distinct compartments, a dorsal one, named the dermomyotome, and a ventral one, called the sclerotome (Parker et al., 2003). The somitic origin of MCs in the limb was first demonstrated in the avian model by performing somite grafting experiments in between chicken and quail embryos (Beresford, 1983; Chevallier et al., 1977; Hayashi and Ozawa, 1995; Jacob et al., 1979; Lance-Jones, 1988; Newman et al., 1981; Ordahl and Le Douarin, 1992; Schramm and Solursh, 1990). Similarly, two distinct mesodermal sources of ECs were identified: (1) the splanchnic mesoderm, which gives rise to ECs vascularizing the viscera and forming the double primitive aortic anlage (Noden, 1989); and (2) the somites, which contribute to the entire endothelial network of the body wall and limbs (Noden, 1989; Pardanaud et al., 1996; Pouget et al., 2006; Wilting et al., 1995). In an equivalent mouse-into-chicken grafting system, the mouse somite gives rise to limb muscles (Houzelstein et al., 1999; Sze et al., 1995; Yvernogeau et al., 2012). Similar to the chicken embryo, the vascularization of the body wall, kidney and limbs were shown to stem from mouse PSM-derived ECs *(Ambler et al., 2001; Yvernogeau et al., 2012)*. The transcription factor Pax3 controls myogenic migration process in both avian and mouse species (Bober et al., 1994; Daston et al., 1996; Goulding et al., 1994; Relaix et al., 2003; Williams and Ordahl, 1994). EC determination is dependent of the vascular endothelial growth factor receptor 2 (Vegfr2), which is encoded by the tyrosine kinase receptor Flk1 gene (Dumont et al., 1998; Fong et al., 1995; Sato et al., 1995; Shalaby et al., 1995; Shivdasani et al., 1995; Visvader et al., 1998).

While the dermomyotome compartment produces ECs and MCs, the sclerotome mainly contributes to the vertebrae and associated cartilage, and produces pericytes and vascular smooth muscle cells (vSMCs) of the aorta, limbs and body wall (Pouget et al., 2006; Wasteson et al., 2008; Wiegreffe et al., 2007, 2009). However, the aorta is formed in a specific developmental process. It is in the aorta that hematopoietic stem cells (HSCs) are first detected. HSCs are found in intra-aortic hematopoietic cluster (IAHCs) that directly derive from specialized hemogenic endothelial (HE) cells embedded in the wall of the aorta (Cai et al., 2000; North et al., 1999; North et al., 2002; Okuda et al., 1996; Wang et al., 1996). In the chicken model, the endothelium of the aorta has clearly a dual embryonic origin (Jaffredo et al., 2010; Pardanaud et al., 1996). The primitive aorta, initially formed by ECs derived from the splanchnopleural mesoderm, is progressively replaced by somite-derived ECs during the fusion of the paired aortas (Jaffredo et al., 2010; Pardanaud et al., 1996; Pouget et al., 2006). Importantly, this replacement coincides with the end of the aortic hematopoietic wave and no IAHCs will be formed anymore. Therefore, somite-derived ECs do not contribute to hematopoiesis in chicken embryos (Jaffredo et al., 2010; Pardanaud et al., 1996; Pouget et al., 2006).

Whether somites also contribute to the aorta in the mouse embryo via a similar mechanism remains to be determined. Mouse somites give rise to highly migratory MCs and ECs, which integrate into tissues such as the limb and body wall (Ambler et al., 2001; Yvernogeau et al., 2012). Unexpectedly, some ECs from the bone marrow vasculature were shown to contribute to adult hematopoiesis (Yvernogeau et al., 2019). However, this study would benefit from an efficient reporter mouse line to further target and trace the somite derivatives. Altogether, it is clear that somite-derived ECs exist, but it remains to be demonstrated whether these cells contribute to either non-HE or HE cells within the mouse aorta.

Despite its importance during embryogenesis, few reporter mouse models were generated to trace the PSM derivatives. PSM is characterized by the specific expression of several *T-box* family of transcription factor genes, such as Brachyury (Herrmann et al., 1990) or Tbx6 (Chapman and Papaioannou, 1998). A recent study demonstrated the utility of a Tbx6 inducible mouse model to trace PSM-derived cells in wild-type and in mutant condition (Concepcion et al., 2017). Although the transgenic mouse appeared efficient to label and study the fate of Tbx6-derived cells in a knockout situation (i.e. in Tbx6^*null*^ embryos), the authors made use of a *LacZ* reporter to describe the progeny and the contribution of the PSM-derived cells in a wildtype situation. No characterization was performed, i.e. immuno-histology, therefore precluding any definitive conclusion on the fate of the *LacZ*^+^ cells obtained. Moreover, the efficiency of the mouse model to tag PSM-derived cells in the adult was not tested (Concepcion et al., 2017). To study in details the PSM-derivatives throughout mouse development and in adults, we generated a new Tbx6 tamoxifen-inducible reporter mouse line (Tg(Tbx6_Cre/ERT2)). Here, we made use of a Tbx6^*Tg122*^ construct previously designed to specifically target the PSM without labelling the tail bud (White et al., 2005). After tamoxifen induction, embryos were collected at sequential developmental time points known to be crucial for the aorta remodeling and limb patterning. Whole-mount immunostaining and immuno-histochemistry on cryosections revealed that our new reporter mouse line faithfully traces the PSM derivatives and can be used to study the formation of ECs and MCs from the embryo until adulthood.

## RESULTS

### Efficient labelling of Tbx6-derived cells after 2 rounds of tamoxifen induction

The expression of Tbx6 starts at E7.5 in the primitive streak (White et al., 2005), and becomes restricted to the PSM and tail bud by E9.5 (**Fig. S1A**). Since our goal was to specifically follow the PSM derivatives, we used a previously reported Tbx6 construct (Tbx6^*Tg122*^) where tagged-cell expression is restricted to the PSM and not to the tail bud (White et al., 2005). Using this construct, we generated a new Tbx6 transgenic reporter line by inserting a tamoxifen-inducible Cre (CreERT2) after the cis-acting regulatory region of the Tbx6 enhancer sequence(White et al., 2005). The Tg(Tbx6_Cre/ERT2) mutant lines obtained were then crossed with the homozygous ROSA26-stop-EYFP mutant line (**Fig. 1A**) to generate a Tg(Tbx6_Cre/ERT2)/ROSA-eYFP line that we used in this study.

**Figure 1.**
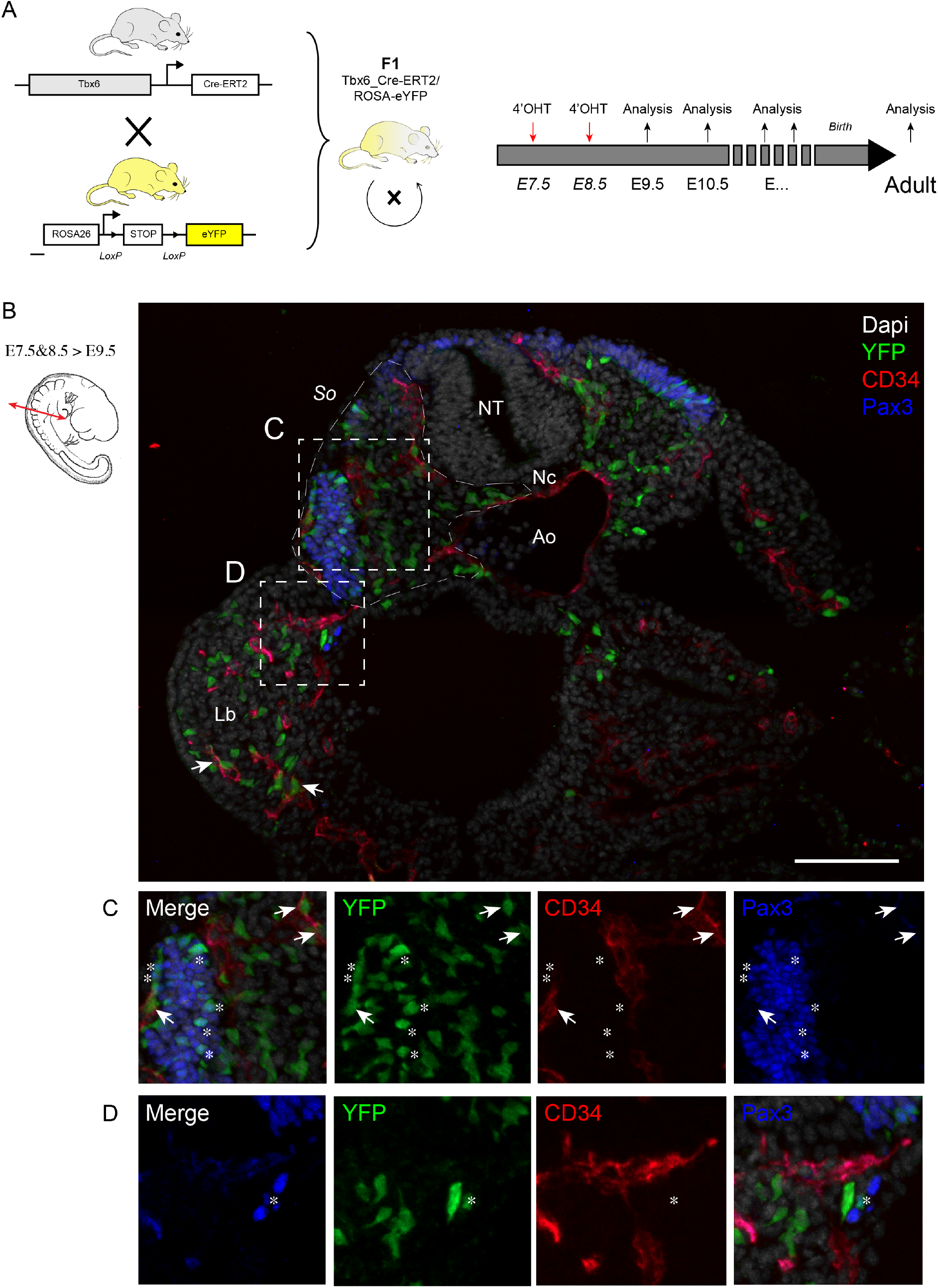
Generation of an inducible Tbx6-cre reporter mouse line to trace the derivatives of the presomitic mesoderm. (A) Experimental procedure scheme. Inducible Tbx6_Cre/ERT2 mice were crossed to the homozygous ROSA26-stop-EYFP mice to generate the Tg(Tbx6_Cre/ERT2)/ROSA-eYFP mutant line. Tg(Tbx6_Cre/ERT2)/ROSA-eYFP mice were then crossed together to generate embryos. Tamoxifen was injected twice, at E7.5 and E8.5, to obtain an efficient cre recombination. Analyses were performed along the developmental course of the embryos and in adults to characterize the progeny of the Tbx6-expressing cells after induction. (B) Representative transversal section of an E9.5 Tg(Tbx6_Cre/ERT2)/ROSA-eYFP embryo immunostained with anti-GFP (green, reflecting Tbx6-derived YFP^+^ cells), anti-CD34 (red, ECs) and anti-PAX3 (blue, myogenic progenitors) antibodies. The section was counterstained with DAPI to visualize nuclei. The dashed squares are shown in C and D as separated single immunostaining pictures. (C) Merge, YFP, CD34 and PAX3 immunostaining pictures showing the contribution of the Tbx6-derived cells to the endothelial (arrows) and myogenic (asterisks) populations at the trunk level. (D) Merge, YFP, CD34 and PAX3 immunostaining pictures showing a Tbx6-derived myogenic progenitor migrating towards the limb bud (asterisk). So, Somite; NT, Neural Tube; Nc, Notochord; Ao, Aorta; Lb, Limb bud. Scale bar: 100μm.

To determine the optimal dose and timing of tamoxifen injections for an efficient induction and tagging of Tbx6-derived cells, we first injected pregnant mice at E8.5 and harvested the embryos the next day (E9.5) for whole-mount immunostaining for CD34 (endothelial maker) and YFP (reflecting the Tbx6-derived cells). Of note, we used GFP antibodies to enhance and unravel the native YFP expression. Only few cells appeared YFP^+^ but were mostly found in the PSM presumptive area, as expected (**Fig. S1B**). No YFP^+^ cells were observed in the forelimb (arrow) and only very rare YFP^+^ cells were found in the anterior somites (**Fig. S1B**, arrowheads). Since the efficiency of the labelling was very low, we tested a double tamoxifen injection of the pregnant mice at both E7.5 and E8.5. This resulted in a more efficient labelling of Tbx6-derived cells (**Fig. S1C**). After whole-mount staining of E9.5 embryos, numerous YFP^+^ cells were indeed observed within the PSM, in all anterior somites and within the forelimb (**Fig. S1C**, arrowheads and arrow, respectively). YFP^+^ cells were also seen to migrate ventrally and to colonize the body wall (**Fig. S1C**, bracket). Importantly, virtually no YFP^+^ cells were observed in the tail bud, as expected from our Tbx6^*Tg122*^ construct (White et al., 2005). At E10.25, the proportion of YFP^+^ cells increased (**Fig. S1D**). These cells were found along the developing embryo, migrating around the neural tube and colonizing both the forelimb and the hindlimb (**Fig. S1D**, arrows). Overall, we determined that two injections of tamoxifen, performed at E7.5 and E8.5, were sufficient to provide an efficient labelling of the Tbx6-derived progeny.

### Lineage tracing of Tbx6-derived cells at E9.5 and E10.5

To characterize the progeny of the Tbx6-derived cells during embryo development, we performed two injections of tamoxifen at E7.5 and E8.5 and analyzed the embryos at E9.5 and E10.5 (**Fig. 1A**). We first performed immunostaining for CD34 (endothelial marker) and Pax3 (somite/myogenic marker) on cryosections obtained from E9.5 Tg(Tbx6_Cre/ERT2)/ROSA-eYFP embryos (**Fig. 1B**). YFP^+^ cells were found dispersed within the forming somite. At the epaxial level, some YFP^+^ cells were present in the dermomyotome, expressing Pax3 (**Fig. 1B,C**, asterisks). Other YFP^+^ cells were observed close to the neural tube or on the top of the dermomyotome, expressing CD34 (**Fig. 1B,C**, arrows). At the hypaxial level, only few YFP^+^Pax3^+^ cells were found migrating and colonizing the limb (**Fig. 1B,D**, asterisk) while some YFP^+^CD34^+^ had already invaded the limb bud (**Fig. 1B**, arrows). This is in accordance with previous reports, showing that the limb is first colonized by PSM-derived ECs and subsequently by PSM-derived myogenic progenitors (Yvernogeau et al., 2012). Of note, some YFP^+^ cells that were neither endothelial nor myogenic cells but most likely mesenchymal-like cells were also present in the limb. Overall, Tbx6-derived cells were found evenly distributed in the embryo providing endothelial and myogenic cells, as previously reported (Ambler et al., 2001; Pardanaud et al., 1996; Pouget et al., 2006; Yvernogeau et al., 2012).

Analyses performed at E10.5 revealed that the proportion of Tbx6-derived cells found in the embryo has increased (**Fig. 2A**). At the hypaxial level, numerous YFP^+^Pax3^+^ myogenic progenitors have invaded the limb or migrated ventrally in the body wall (**Fig. 2A-E**, white and yellow arrowheads, respectively). Sparsely, YFP^+^CD34^+^ ECs that contributed to the vascularization could be identified (**Fig. 2A-E**, red arrows). At the epaxial level, Tbx6-derived cells have contributed to the formation of the perineural vascular plexus (PNVP) (**Fig. 2A and F-I**, arrowheads). In the dorsal aortic region, Tbx6-derived cells contributed to the sclerotome and were present all around and in close contact to the endothelium of the aorta (**Fig. 2J-M**, asterisks). We observed that some Tbx6-derived cells could contribute to the endothelium and to the smooth muscle cells forming the dorsal aorta (**Fig. 2N-Q**, arrowheads and arrows, respectively). However, and very importantly, no contribution of Tbx6-derived cells to IAHCs was found (**Fig. 2J-M**, arrowheads).

**Figure 2.**
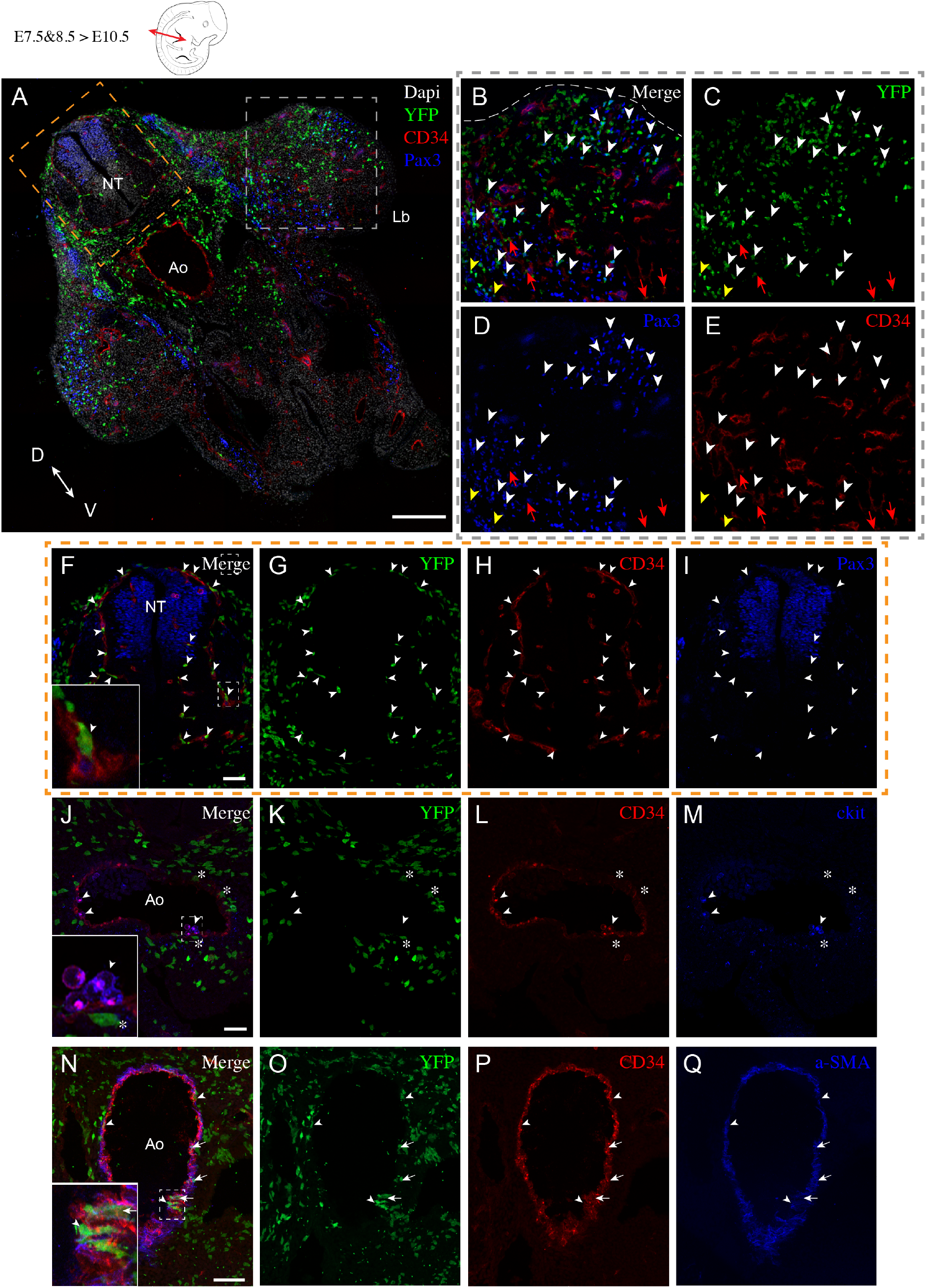
Lineage tracing shows the contribution of Tbx6-derived cells to the endothelial and myogenic lineages of the trunk and limbs at E10.5. (A) Representative E10.5 transversal section of a Tg(Tbx6_Cre/ERT2)/ROSA-eYFP embryo, at the forelimb level, immunostained with anti-GFP (green), anti-CD34 (red), anti-PAX3 (blue) antibodies and counterstained with DAPI. Grey dashed square is shown as separated immunostaining pictures in B-E. Orange dashed square is shown as separated immunostaining pictures in F-I. (B-E) Analysis at the limb level. Merge (B), YFP (C), PAX3 (D) and CD34 (E) immunostaining pictures showing the contribution of Tbx6-derived cells to (i) the myogenic cells colonizing the limb bud (white arrowheads) and the body wall (yellow arrowheads), (ii) the vascularization of the limb (red arrows). (F-I) Analysis at the neural tube level. Merge (F), YFP (G), CD34 (H) and PAX3 (I) immunostaining pictures showing the contribution of Tbx6-derived cells to the perineural vascular plexus (PNVP) (arrowheads). Higher magnification of the dashed square in (F) is shown enlarged on the bottom left corner of panel F. (J-M) Dorsal aorta level from the adjacent cryosection shown in (F-I). Merge (J), YFP (K), CD34 (L) and c-kit (M) immunostaining pictures showing Tbx6-derived cells close to the endothelium of the aorta (asterisks) but not in the intra-aortic hematopoietic cluster cells (arrowheads). Higher magnification of the dashed square in (J) is shown enlarged on the bottom left corner of panel J. (N-Q) Dorsal aorta level from an adjacent cryosection. Merge (N), YFP (O), CD34 (P) and a-SMA (Q) immunostaining pictures showing Tbx6-derived cells contributing to the endothelium (arrowheads) and to the smooth muscle (arrows) of the aorta. Higher magnification of the dashed square in (N) is shown enlarged on the bottom left corner of panel N. Ao, Aorta; NT, Neural tube; Lb, Limb bud; D, Dorsal, V, ventral. Scale bars: 200μm in A-E; 50μm in F-Q.

### Tbx6-derived cells give rise to all PSM derivatives throughout development

To determine whether our Tg(Tbx6_Cre/ERT2)/ROSA-eYFP line faithfully labelled all PSM derivatives after mid-gestation, we performed analyses on E11.5 embryos (**Fig. 3**). On transversal cryosections obtained at the forelimb level, YFP^+^ cells were observed within the differentiated somite, from the dermomyotome area to the sclerotome area (**Fig. 3A**). As expected from the previous data obtained at E9.5 and E10.5 (**Fig. 1 and 2**), YFP^+^ cells were also present in the forelimb. At the epaxial level, YFP^+^ cells had now virtually formed the entire PNVP (**Fig. 3A** and **Fig. S2A-D**, arrowheads). Only rare ECs did not express YFP within the neural tube (**Fig. S2A-D**, asterisks). At the level of the aorta, very few YFP^+^ cells contributed to the endothelium (**Fig. 3A** and **Fig. S2E-H**, arrowheads) while others were integrated into the smooth muscle layer/tunica of the aorta (**Fig. S2I-L**, arrowheads). This is consistent with our previous observations at earlier developmental time points (**Fig. 1 and 2**) and in accordance with previous reports (Pouget et al., 2006; Pouget et al., 2008; Wiegreffe et al., 2007; Yvernogeau et al., 2012). At the hypaxial level, myogenic cells that have emigrated from the somite have split into two separated muscle masses, ventral and dorsal (**Fig. 3A**, brackets), both showing contribution of YFP^+^ cells (**Fig. 3A** and **Fig. S2M-P**, arrowheads). YFP^+^CD34^+^ ECs also contributed to the vascularization of the limb until its most distal part (**Fig. 3A**, arrowheads and **Fig. S2Q-T**, arrows). Of note, immunostaining with EC surface marker (such as CD34) does not overlap with the YFP since our Tbx6-ERT2 construct allows YFP expression in the cytoplasm of the cells. To ascertain the presence of YFP^+^ ECs within the trunk and limbs of the Tg(Tbx6_Cre/ERT2)/ROSA-eYFP embryos, we performed flow cytometry analysis (**Fig. 3B**). This also allowed us to quantify the YFP^+^ and the EC populations in the dissected trunk and limbs. We found that the trunk and limb contained an average of 8.2% and 5% of YFP^+^ cells, respectively (**Fig. 3C,D** and **Table S1**). Within the YFP^+^ population, only a small fraction contributed to the endothelial lineage, i.e. 0.7% in the trunk and 0.4% in the limb (**Fig. 3C,D** and **Table S1**). On the other hand, Tbx6-derived ECs (i.e. YFP^+^) represented 4.3% and 3.6% of the total endothelial population within the trunk and limb, respectively (**Fig. 3C,D** and **Table S1**).

**Figure 3 (see also Figure S2).**
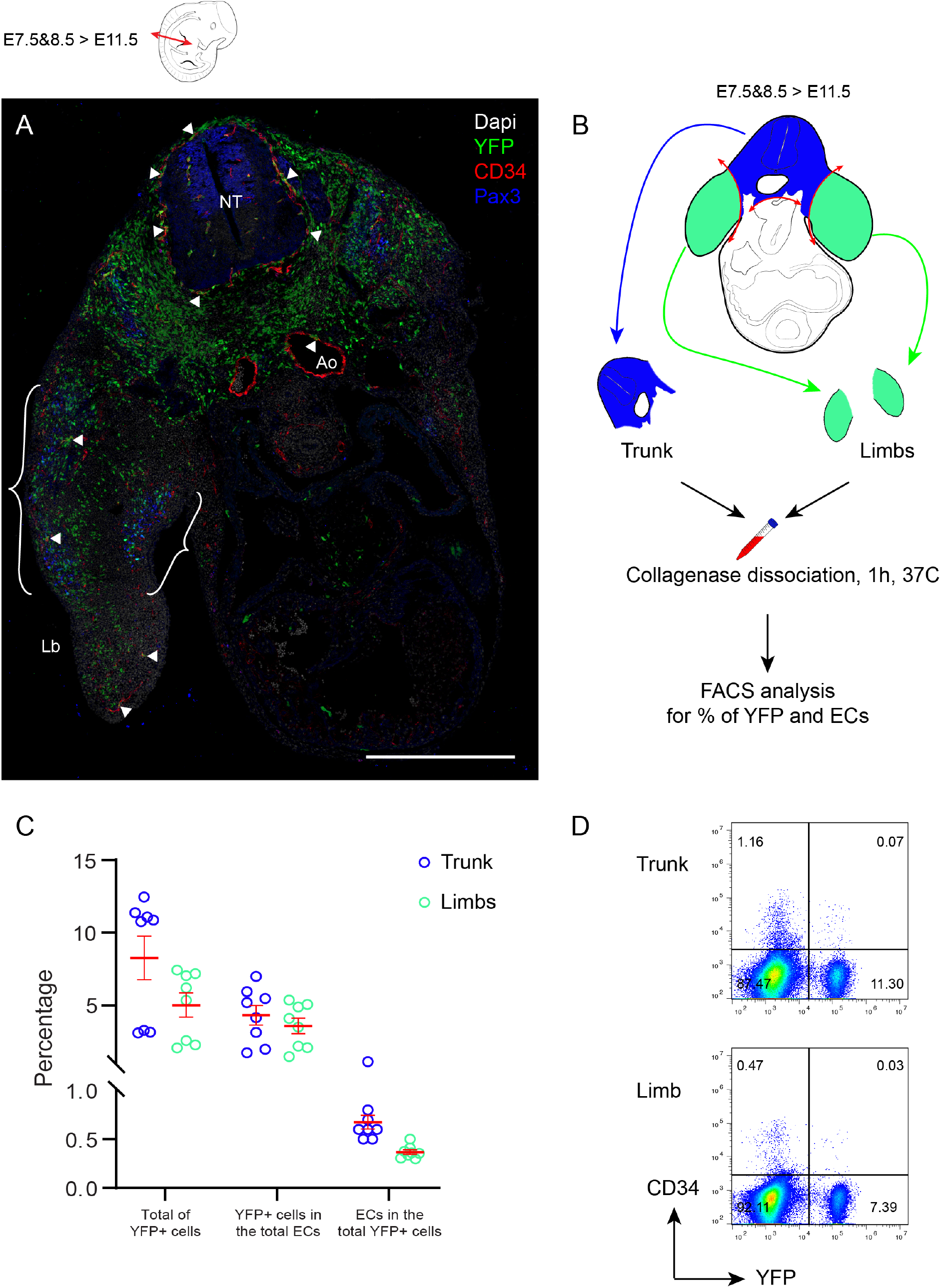
Lineage tracing shows the contribution of Tbx6-derived cells to both the limb muscle masses and the endothelial population at E11.5. (A) Representative E11.5 transversal section of a Tg(Tbx6_Cre/ERT2)/ROSA-eYFP embryo, at the forelimb level, immunostained for YFP (green), CD34 (red), PAX3 (blue) and counterstained with DAPI. (B) Experimental procedure used to isolate and dissociate the limbs and trunks of E11.5 Tg(Tbx6_Cre/ERT2)/ROSA-eYFP embryos for flow cytometry analysis. (C) Graph representing the percentages of (i) YFP^+^ cells in the trunk or limb, (ii) YFP^+^ cells in the endothelial population and (iii) ECs within the YFP^+^ population. One circle represents one embryo. Eight embryos from two different litters were used. Data are represented as mean±SEM. (D) Representative FACS plots showing the percentages of YFP^+^ cells and CD34^+^ cells (i.e. ECs) in the trunk and limbs. Ao, Aorta; NT, Neural tube; Lb, Limb Scale bar: 500μm.

To further explore the expression pattern of the Tbx6-derived cells throughout development, we analyzed Tg(Tbx6_Cre/ERT2)/ROSA-eYFP embryos at E14.5 (**Fig. 4** and **Fig. S3**). Immunostainings performed on transversal cryosection at the limb level revealed the global contribution of Tbx6-derived cells (**Fig. 4A**). At the epaxial level, in the dorsal part of the embryo, YFP^+^ cells contributed to muscles and to the vasculature (**Fig. 4A, B** and **Fig. S3A-D**, arrows and arrowheads, respectively). Of note, some YFP^+^ cells started to condense to form the vertebral skeleton (**Fig. 4B** and **Fig. S3A-D**, encircled area). Similar, in the ventral part of the embryo, other YFP^+^ cells contributed to the formation of the ribs (**Fig. 4A**, curly bracket). This is in accordance with the presence of YFP^+^ cells within the sclerotome (**Figs. 2A** and **3A**), the somite compartment which provides cells that will form bones and cartilages of the trunk.

At the hypaxial level, YFP^+^ cells formed the differentiated muscles and participated to the vasculature (**Fig. 4A, C** and **Fig. S3E-H**, arrows and arrowheads, respectively). We also performed flow cytometry analysis on cells obtained after the trunk and limbs were dissected and dissociated to assess for the presence of YFP^+^ cells and CD34^+^ ECs. We found that E14.5 trunk and limb contained an average of 8.1% and 3.9% of YFP^+^ cells, respectively (**Fig. 4D,E** and **Table S1**), a similar proportion as previously observed at E11.5 (**Fig. 3C,D** and **Table S1**). Within the YFP^+^ population, the contribution to the endothelial lineage was 1.3% in the trunk and 1.1% in the limb (**Fig. 4D 4,E** and **Table S1**). Compared to the data obtained at E11.5, these populations have significantly increased of 18% and 27.5% (p<0.0002 and p<0.0006, for trunk and limb, respectively) suggesting that YFP^+^ ECs have proliferated. Finally, Tbx6-derived ECs (i.e. YFP^+^) contributed to 7.6% and 3.4% of the total endothelial population within the trunk and limbs, respectively (**Fig. 3C,D** and **Table S1**), showing an increase of this population in the trunk and a similar proportion in the limbs when compared to data obtained at E11.5.

**Figure 4 (see also Figure S3).**
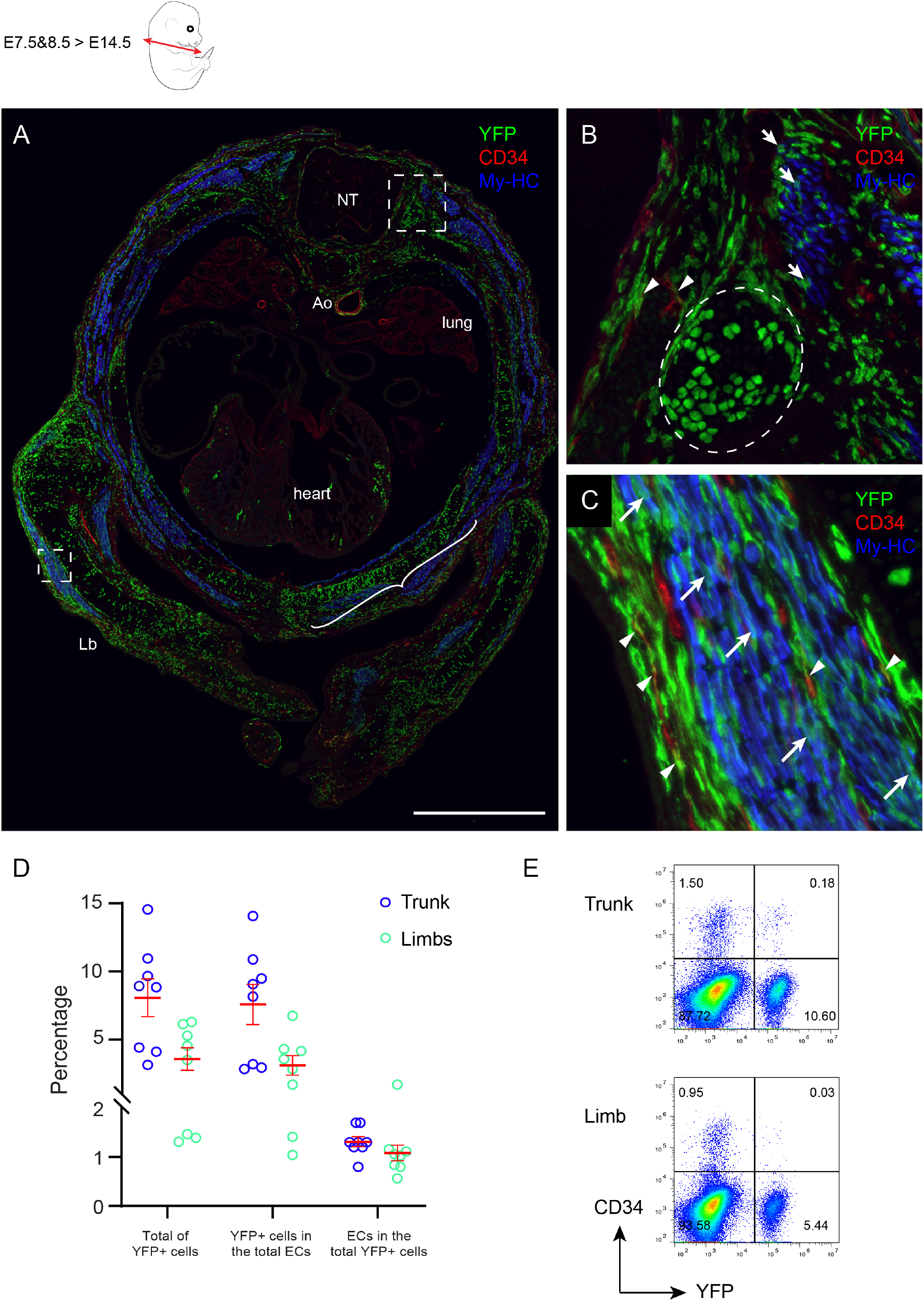
Lineage tracing shows the contribution of Tbx6-derived cells to vertebrae and limb muscles, and an increased contribution to the endothelial population at E14.5. (A) Representative E14.5 transversal section of a Tg(Tbx6_Cre/ERT2)/ROSA-eYFP embryo, at the forelimb level, immunostained for YFP (green), CD34 (red) and MyHC (blue). Dashed squares are shown enlarged in B and C. (B) Enlarged view of the trunk area indicated in (A) showing Tbx6-derived cells contributing to the cartilage (encircled area), the vascularization (arrowheads) and muscles (arrows). (C) Enlarged view of the limb area indicated in (A) showing Tbx6-derived cells contributing to the vascularization (arrowheads) and to the muscles (arrows). (D) Graph representing the percentages of (i) YFP^+^ cells in the trunk or limb, (ii) YFP^+^ cells in the endothelial population and (iii) ECs within the YFP^+^ population. One circle represents one embryo. Eight embryos from two different litters were used. Data are represented as mean±SEM. (E) Representative FACS plots showing the percentages of YFP^+^ cells and CD34^+^ cells (i.e. ECs) in the trunk and limbs. Ao, Aorta; NT, Neural tube; Lb, Limb bud. Scale bar: 1mm.

### The Tg(Tbx6_Cre/ERT2)/ROSA-eYFP line is suitable to label PSM-derived cells until late embryonic stages and in adults

To evaluate whether our Tg(Tbx6_Cre/ERT2)/ROSA-eYFP line could be used to study late developmental processes, we analyzed embryos at E16.5 (**Fig. 5** and **Fig. S4**). Immunostainings performed on transversal cryosections of the hindlimb revealed that YFP^+^ cells were distributed throughout the entire tissue (**Fig. 5A**). YFP^+^ cells had colonized the bone marrow, participating to the vascularization (**Fig. 5A-E**). Other YFP^+^ cells, but CD31^-^CD34^-^, were also present. They most likely represent mesenchymal-like cells since no contribution to the hematopoietic lineage was observed. Indeed, no YFP^+^ cells expressing c-kit were ever found. The muscles of the hindlimb were virtually all YFP^+^ (**Fig. 5A, F-I** and **Fig. S4A-D**), demonstrating that our Tg(Tbx6_Cre/ERT2)/ROSA-eYFP line can be used to efficiently label the muscle progeny of the PSM. Importantly, we observed the same contribution by the YFP^+^ cells within the forelimb (**Fig. S4A-D**). In both transversal or longitudinal sections, the muscles were almost completely YFP^+^, from the proximal (**Fig. S4A,B**) to the distal part of the limb (**Fig. S4C,D**).

**Figure 5 (see also Figure S4).**
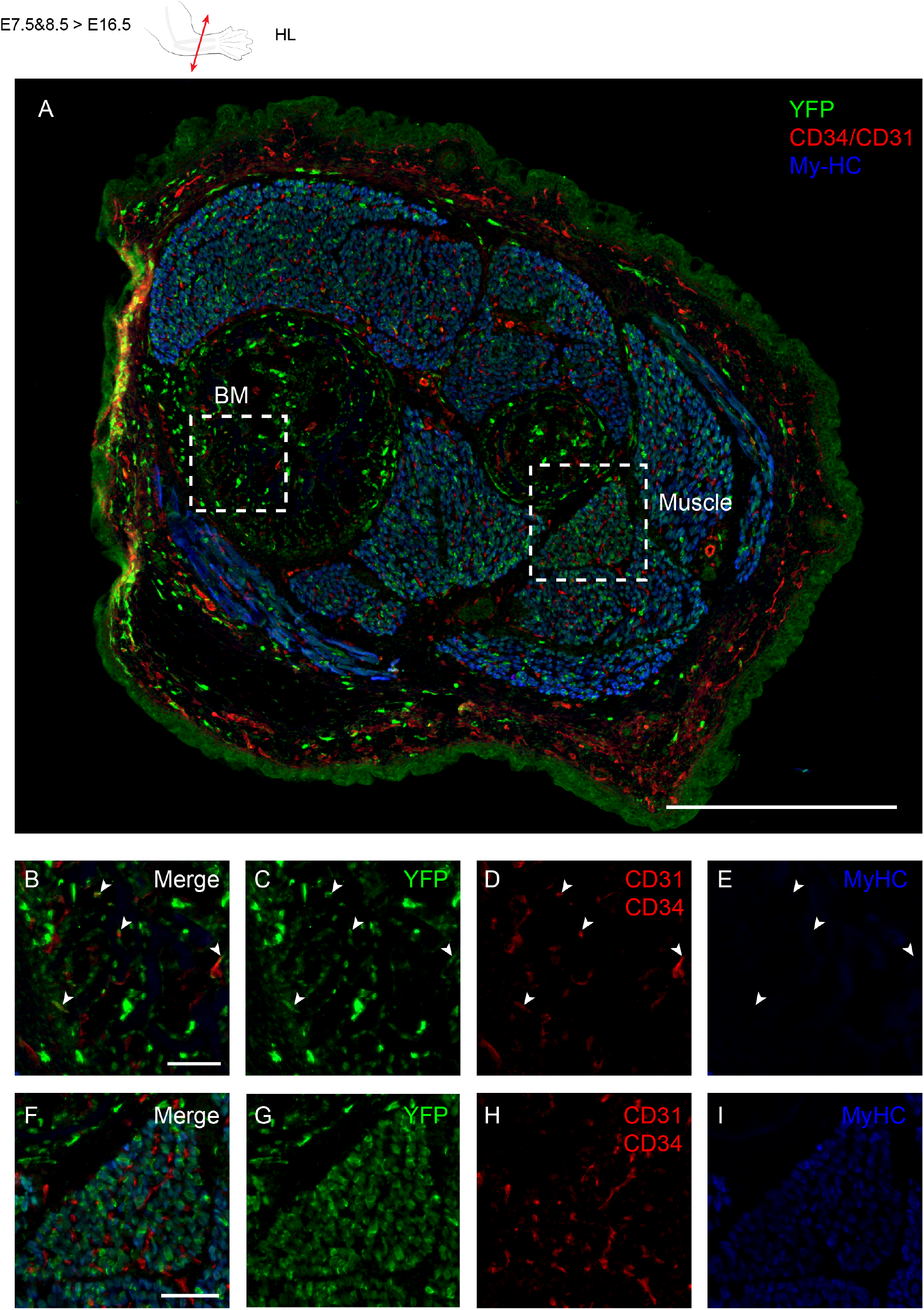
Lineage tracing shows the contribution of Tbx6-derived cells to the limb musculature and vascularization at E16.5. (A) Representative E16.5 transversal section of a Tg(Tbx6_Cre/ERT2)/ROSA-eYFP hindlimb, immunostained for YFP (green), CD34 and CD31 (red) and MyHC (blue). Dashed squares are shown enlarged in B-E and F-I. (B-E) Enlarged view of the bone marrow area indicated in (A) showing Tbx6-derived cells contributing to the vascularization (arrowheads). (F-I) Enlarged view of the muscle area indicated in (A) showing Tbx6-derived cells contributing to the entire muscle. Scale bars: 500μm in A; 50μm in B-I.

Finally, we analyzed the contribution of the Tbx6-derived cells in adult tissues (**Fig. 6** and **Fig. S5**). Two-month-old adult mouse legs were isolated and then immunostained for CD31/CD34 endothelial markers (combined antibodies) and for laminin. The entire musculature of the legs was YFP^+^, therefore originating from PSM-derived cells (**Fig. 6A** and **Fig. S5A-D**). The vascularization was also partially derived from Tbx6-derived cells (**Fig. 6B-E** and **Fig. S5A-D**, arrowheads). Some YFP^+^ nuclei were observed underneath the basal lamina of the muscle fibers (**Fig. 6B-D** and **Fig. S5A-D**, arrows), in a location where the muscle stem cells (i.e. the satellite cells) are normally found. This is in accordance with previous reports showing that somites are the source of adult limb muscle satellite cells (Gros et al., 2005; Relaix et al., 2005). Overall, our Tg(Tbx6_Cre/ERT2)/ROSA-eYFP line is a reliable model to study and trace PSM-derived cells throughout development until adulthood.

**Figure 6 (see also Figure S5).**
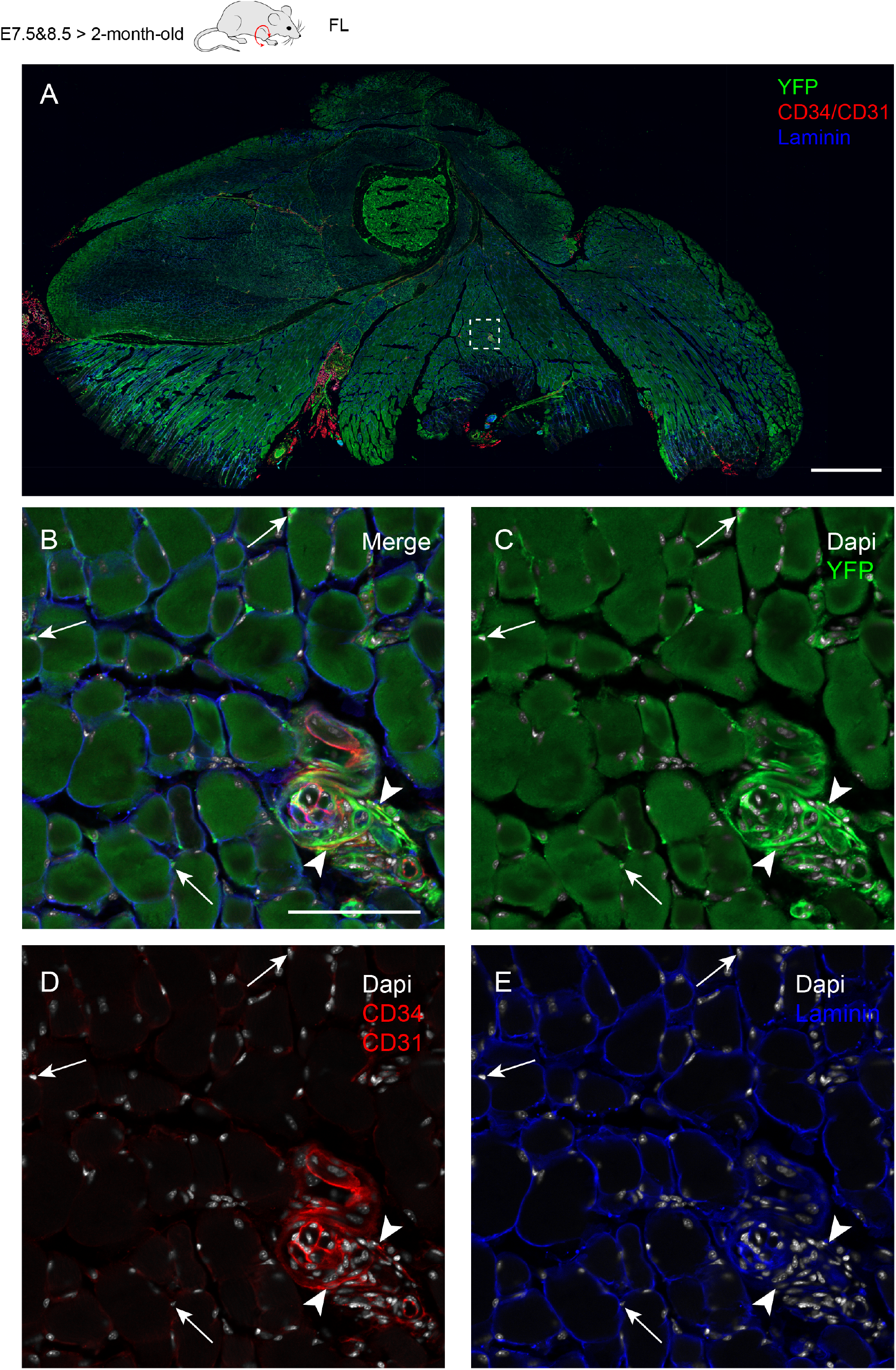
Long-term lineage tracing of Tbx6-derived cells after induction at E7.5 and E8.5 shows muscle contribution in adult mice. (A) Transversal section of the forelimb isolated from a two-month-old Tg(Tbx6_Cre/ERT2)/ROSA-eYFP mouse, immunostained for YFP (green), CD34 and CD3l (red) and laminin (blue). Dashed square is shown enlarged in B-E. (B-E) Enlarged view of the muscle area indicated in (A) showing Tbx6-derived cells contributing to the vascularization (arrowheads). Some YFP^+^ nuclei localized underneath the basal lamina of the muscle fibers (arrows). Scale bars: 1.000μm in A; 100μm in B-E.

## DISCUSSION

The PSM is an unsegmented mesoderm that progressively segments to form somites, round structures found on each side of the neural tube (Parker et al., 2003). Somites are at the foundation of the musculature of the trunk and limbs but also provides the vasculature, cartilage, bones and smooth muscle cells of the arteries. Despite its importance for embryogenesis, there is still a lack of efficient reporter mouse line to trace the PSM derivatives. In the present study, we generated a new inducible transgenic reporter mouse line that we named Tg(Tbx6_Cre/ERT2)/ROSA-eYFP, which allows to permanently label and trace the PSM derivatives until adulthood. After optimizing the induction procedure and verifying the label efficiency, we found that Tbx6-derived cells quickly (< 24hours) contributed to the endothelial vascular network around the neural tube, i.e. the PNVP, as well as the forming limb bud, in a same time frame as previously reported in different species and contexts (Ambler et al., 2001; Pouget et al., 2006; Yvernogeau et al., 2012). The Tbx6 derivatives were also found within the developing somite, in the dorsal (dermomyotome) and ventral (sclerotome) parts. At this stage, the first myogenic progenitors coming from the somites also started to migrate and colonize the limb while they expressed the transcription factor Pax3 (Yvernogeau et al., 2012). Later, Tbx6-derived cells contributed not only to the vasculature of the limbs but also to the dorsal and ventral muscle masses. Tbx6-derived myogenic cells were also found migrating within the body wall. Of note, we observed that only few Tbx6-derived cells contributed to the endothelial lineage. Nevertheless, with the embryonic development progressing, this proportion significantly increased demonstrating that either this population proliferated with time, or that the specific Tbx6-derived endothelial population is ‘selected’ during limb vascular remodeling. However, further investigation will be needed to verify these hypotheses. After the limb muscle splitting process, we observed that Tbx6-derived cells contributed to all the differentiated muscles. It seems unlikely that Tbx6-derived cells constitute the entirety of the muscle, as one shall consider that myogenic progenitors fuse to form the muscle fibers, which might explain this unexpected contribution. Further investigation will be required to determine the proportion of Tbx6-derived myogenic cells that actually contributes to the limb muscles. Besides the musculature of the limb, we identified some Tbx6-derived cells invading the bone marrow of the late embryos (>E16) and contributing to its vascularization, as it was previously reported (Yvernogeau et al., 2019). Finally, our Tg(Tbx6_Cre/ERT2)/ROSA-eYFP line appears as a suitable model to trace at long-term Tbx6-derived cells. Indeed, the vasculature and musculature of the adult limbs were derived from Tbx6-derived cells. We identified YFP^+^ nuclei also located underneath the basal lamina of the muscle fibers, showing that our Tg(Tbx6_Cre/ERT2)/ROSA-eYFP is suitable to tag and trace muscle stem cells at long-term.

In the present study, we confirmed the existence of somite-derived ECs that contribute to the endothelium of the aorta, but never to IAHCs. IAHCs have been previously observed near the somites, within the inters-omitic vessels and dorso-lateral anastomotic vessels (Yzaguirre and Speck, 2016). This may indicate the presence of somite-derived HE cells that might have the potential to generate HSCs. However, the authors could not exclude that these IAHCs migrated to these finer vessels through the blood circulation rather than being produced there *de novo*. In support of this hypothesis, the presence of somite-derived aortic ECs has also been shown in zebrafish, in which these cells arise from a specific somite compartment that the authors characterized as the ‘endotome’ (Nguyen et al., 2014). These cells do not directly produce HE cells but rather indirectly contribute to HSC generation through chemokine signaling to HE cells within the dorsal aorta that are derived from the lateral mesoderm. Qiu *et al*. identified somite-derived HE cells, which gives rise to multipotent and self-renewing HSCs but these findings remain to be confirmed (Qiu et al., 2016). Here, the data obtained with our Tg(Tbx6_Cre/ERT2)/ROSA-eYFP mouse model support the notion that PSM-derived cells do not contribute to HE cells within the mouse aorta.Generation of an inducible Tbx6-cre reporter mouse line to trace the derivatives of the presomitic mesoderm.

## MATERIALS AND METHODS

### Mice

#### Tbx6_Cre/ERT2 mouse line generation

The Tg(Tbx6_Cre/ERT2) transgenic mouse line was generated by insertion of a tamoxifen-inducible Cre (CreERT2) after the cis-acting regulatory region of Tbx6 sequence (a 2.3kb fragment driving reporter gene expression in the presomitic mesoderm (White et al., 2005) under minimal promoter control (hsp68). The initial plasmid construct was kindly provided by Pr. D.L. Chapman. After clone selection, plasmid construct was injected into one cell-stage mouse embryos. Several Tg(Tbx6_Cre/ERT2) mutant lines were obtained giving rise to the same overall expression patterns. Selected Tg(Tbx6_Cre/ERT2) lines were then crossed with the homozygous ROSA26-stop-EYFP mutant mice, which have a *loxP* flanked STOP sequence followed by enhanced yellow fluorescent protein gene (EYFP) inserted into the *Gt(ROSA)26Sor* locus (Srinivas et al., 2001). The resulting double transgenic mice [Tg(Tbx6e_Cre/ERT2)/ROSA-eYFP] were then crossed together and use to generate embryos for all follow-up experimental procedures.

#### Induction with 4-hydroxytamoxifen (4’OHT)

4-Hydroxytamoxifen (Sigma H-7904) was dissolved in sunflower oil (Sigma S-5007) by sonication without overheating to a final concentration of 5mg/ml. The 4’OHT stock solution was stored and protected from the light at −20°C. Intraperitoneal injections of 2mg 4’OHT were administered to the pregnant female mice at E7.5 and E8.5. For the timing of 4’OHT injections, the day of vaginal plug was considered as day 0. All animals were housed according to institutional guidelines, and procedures were performed in compliance with Standards for Care and Use of Laboratory Animals with approval from the Hubrecht Institute ethical review board. All animal experiments were approved by the Animal Experimentation Committee (DEC) of the Royal Netherlands Academy of Arts and Sciences.

#### Mouse embryo generation and collection

Mouse embryos were generated from timed matings. Mouse embryos were collected at embryonic day (E)9.5, E10.5, E11,5, E14.5 and E16.5. For E9.5 and E10.5, embryos were precisely staged based on the number of somite pairs (sp), 20-25 and 30-36sp, respectively.

At the expected/desired developmental stages, embryos were removed from the uterus and yolk sac, and washed in PBS/FCS (PBS supplemented with 10% fetal calf serum). Head was cut off with needles and a piece of the tissue was used to genotype the embryo. Blood was flushed out of the aorta by using a borosilicate needle containing PBS/FCS. Embryos/tissues (i.e. limbs for late embryos > E14) were then fixed in a 4% paraformaldehyde (PBS/PFA) solution during 30 minutes at 4°C.

#### Tissue preparation for whole-mount stainings

After fixation, embryos/tissues were washed three times with PBS before dehydration-rehydration step in methanol. Embryos were then blocked in a PBS-MT (1% skim milk, 0.4% TritonX-100, 0.2% BSA (bovine serum albumin [Sigma]), 0.1% goat serum) solution for 1 hour at 4°C. Embryos/tissues were stained with (i) biotin anti-CD34 antibody (ebiosciences, clone RAM34) to stain the endothelial cells, and (ii) purified anti-GFP antibody (Abcam, ab290), which cross-reacts with the native YFP to stain the progeny of the Tbx6-derived cells. Alexa Fluor 555-Streptavidin and Alexa Fluor 488 goat anti-rabbit antibodies (both from Life Technologies) were used to reveal CD34 and GFP antibodies, respectively.

#### Tissue preparation for immuno-histology

After fixation, embryos/tissues were washed three times with PBS, then cryoprotected in a PBS/15% sucrose solution. Finally, embryos/tissues were embedded in gelatin (7.5%) dissolved in the PBS/15% sucrose solution, then frozen in liquid nitrogen. Sections (10μm) were made using a Leica cryostat. After re-hydratation and gelatin removal in warmed PBS solution, sections were blocked with a PBS/10% FCS/1% BSA/10% goat serum solution (Blocking Solution, BS) for 1 hour. Primary antibodies (see Table 1 below) were incubated in BS overnight at 4°C. After several and extensive washes in PBS (at least 3 times), secondary antibodies were incubated in BS at room temperature for 1h30. Finally, sections were counterstained with DAPI to visualize nuclei.

**Table 1.**
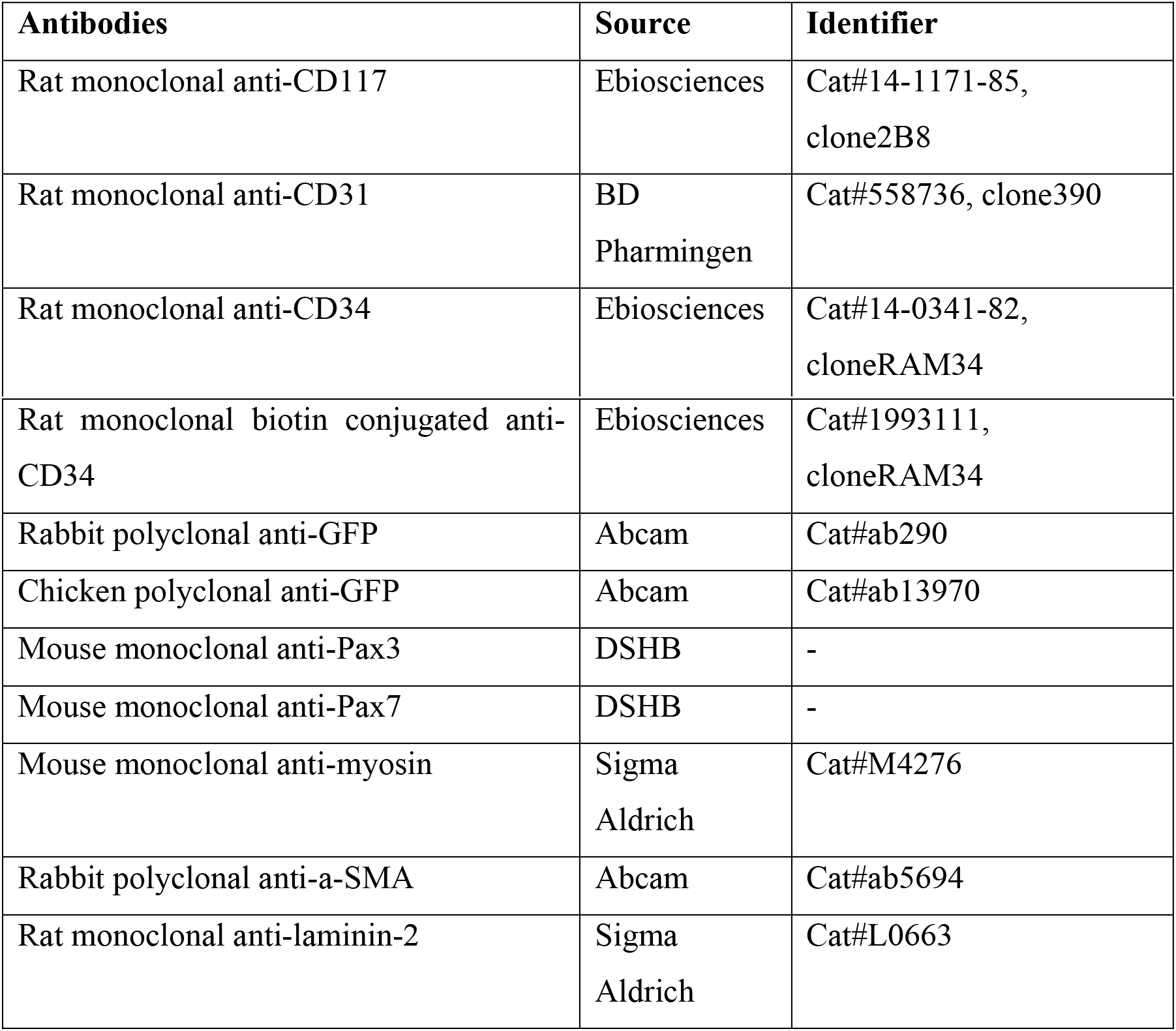
List of the primary antibodies used for immuno-histology.

| **Antibodies** | **Source** | **Identifier** |
| --- | --- | --- |
| Rat monoclonal anti-CD117 | Ebiosciences | Cat#14-1171-85,<br>clone2B8 |
| Rat monoclonal anti-CD31 | BD<br>Pharmingen | Cat#558736, clone390 |
| Rat monoclonal anti-CD34 | Ebiosciences | Cat#14-0341-82,<br>cloneRAM34 |
| Rat monoclonal biotin conjugated anti-CD34 | Ebiosciences | Cat#1993111,<br>cloneRAM34 |
| Rabbit polyclonal anti-GFP | Abcam | Cat#ab290 |
| Chicken polyclonal anti-GFP | Abcam | Cat#ab13970 |
| Mouse monoclonal anti-Pax3 | DSHB | - |
| Mouse monoclonal anti-Pax7 | DSHB | - |
| Mouse monoclonal anti-myosin | Sigma<br>Aldrich | Cat#M4276 |
| Rabbit polyclonal anti-α-SMA | Abcam | Cat#ab5694 |
| Rat monoclonal anti-laminin-2 | Sigma<br>Aldrich | Cat#L0663 |

### Whole adult leg preparation for immunohistochemistry on cryosections

Two-month-old Tg(Tbx6_Cre/ERT2)/ROSA-eYFP mouse legs were dissected and skin was removed before transferred into cold 4% PFA solution and fixed overnight at 4°C under rotation. Next, legs were washed several times in PBS solution then transferred into 10X EDTA solution (pH7.5-8) and kept at 4°C under rotation for 2/3 weeks. EDTA solution was refreshed every two days. Then, samples were dehydrated in a 20% sucrose/1% polyvinylpyrrolidone/PBS solution for at least 3 days (solution being refreshed every day) at 4°C under rotation. Finally, samples were embedded in PBS containing 15% sucrose, 8% gelatin and 1% polyvinylpyrrolidone (overnight at 37°C, under rotation) and frozen in liquid nitrogen. 12 to 14μm section were done using a Leica cryostat.

### Confocal microscopy image acquisition

All cryosections and whole embryos were imaged with a Zeiss LSM900 confocal microscope with a 20xPlanApo dry objective.

### *In situ* hybridization

*EcoRI* and *XhoI* restriction sites were added to the primers for selective ligation into the PCS2 plasmid vector. Plasmid was linearized and riboprobe transcribed from linearized template in the presence of digoxigenin-11-UTP. E9.5 mouse embryos were fixed in 4% PFA solution overnight, and dehydrated stepwise into methanol before storage at −20°C. Embryos were rehydrated stepwise in PBST, and then incubated for 15 minutes with Proteinase K (20μg/ml) followed by 20 minutes post-fixation in 4% PFA. Embryos were prehybridized in Hyb-buffer containing Dextran Sulfate (10X final concentration), Denhart (1X final concentration) and CHAPS (1%) solution for at least 1 hour at 70°C. Riboprobes were diluted 1/200 in Hyb-buffer supplemented with transfer RNA and heparin, and embryos were incubated in probe solution overnight at 70°C. Following probe removal, embryos were extensively washed in MABT solution (pH7.5). Embryos were blocked in PBST with BSA and Sheep Serum for at least 1 hour, and incubated with anti-digoxigenin AP antibody overnight at 4°C. Following antibody incubation embryos were washed extensively in PBST and transferred into TBST. Probe detection was carried out using NBT/BCIP. Embryos were fixed in 4% PFA and then either washed stepwise to PBS/Glycerol (80%) for whole embryo pictures or processed for freezing/cryosectioning (for transversal section pictures).

The following sequences were used to synthetize the Tbx6 probe:

5’ - GGA*GAATTC*GCTGTGGGGACAGAGATGAT-3’ *EcoRI site*
5’ - AGG*CTCGAG*ATCCCGCTCCCTCTTACAGT-3’ *XhoI site*

Expected probe size: 552bp

### Flow cytometry cell analysis

E11.5 and E14.5 Tg(Tbx6_Cre/ERT2)/ROSA-eYFP tamoxifen-induced embryos were used. Trunks and limbs were dissected (see ***Fig. 3B***) and dissociated by collagenase treatment (0.12% w/v, type I, Sigma) for 1 hour or 1h30 (for E14.5 tissues) at 37°C. After dissociation, cells were washed and stained using CD31-APC (clone 390, ebioscience, cat17-0311-82) endothelial antibodies for 30 minutes at 4°C. After washing the antibodies, DAPI was added to exclude dead cell. Analysis was performed using a CytoFLEX S (Beckman Coulter).

## AUTHOR CONTRIBUTIONS

L.Y. conceived ideas, designed the research, performed experiments, analyzed the data and wrote the manuscript. A.K. helped for embryo tissue preparations and commented on the manuscript. C.v.R. generated the Tg(Tbx6_Cre/ERT2) transgenic mouse line. C.R. supervised the study, analyzed the results, wrote the manuscript and acquired funding.

