## Supplemental Fig S1 for "Generation of a new Tbx6-inducible reporter mouse line to trace presomitic mesoderm derivatives throughout development and in adults"

**Supplemental Figure S1**

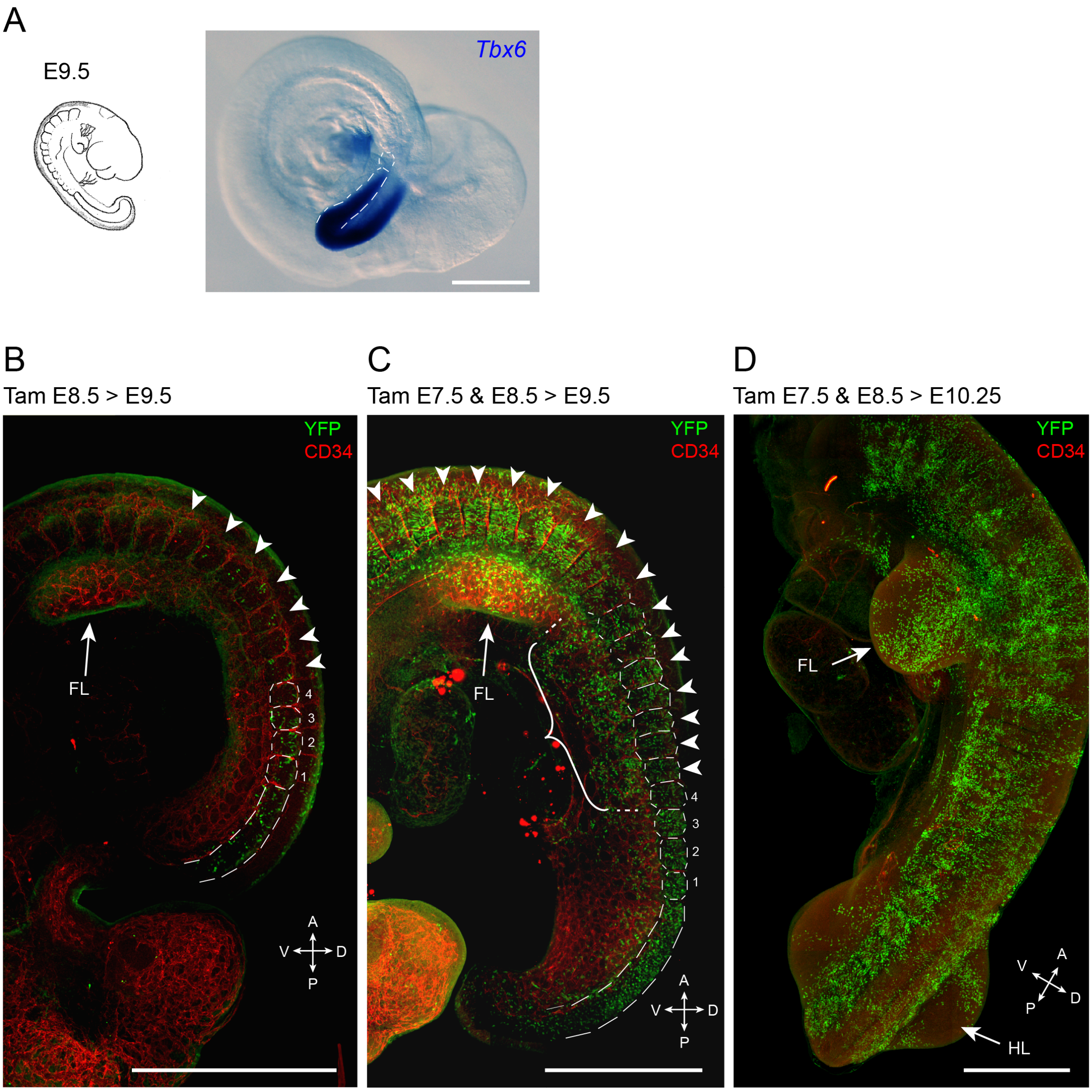


**The injection of 2mg tamoxifen at both E7.5 and E8.5 efficiently induces YFP expression to label and trace the Tbx6-derived cell progeny.**

(A) Whole-mount *in situ* hybridization for Tbx6 on an E9.5 wild-type mouse embryo. At this stage, the presomitic mesoderm is the only structure expressing TBX6. (B) Whole-mount immunostaining for CD34 (red) and YFP (green) on an E9.5 Tg(Tbx6_Cre/ERT2)/ROSA-eYFP mouse embryo, which had received one tamoxifen injection of 2mg at E8.5. Of note, only few $YFP^+$ cells (i.e. $Tbx6^+$-derived cells) are labelled within the PSM and only rare $YFP^+$ cells are found in the anterior developing somites (arrowheads). (C) Whole-mount immunostaining for CD34 and YFP on an E9.5 Tg(Tbx6_Cre/ERT2)/ROSA-eYFP mouse embryo, which had received one tamoxifen injection of 2mg at E7.5 and another tamoxifen injection of 2mg at E8.5. Of note, the number of YFP$^+$ cells that were induced are far more numerous and specifically found within the anterior somites (arrowheads) and in the developing PSM. YFP$^+$ cells colonized the forelimb (arrow) and started to migrate to colonize the body wall (curly bracket). (D) Whole-mount immunostaining for CD34 and YFP on an E10.25 Tg(Tbx6_Cre/ERT2)/ROSA-eYFP mouse embryo, which have received one tamoxifen injection (2mg) at E7.5 and another tamoxifen injection (2mg) at E8.5. The efficient induction of the Tbx6$^+$ (i.e. PSM-)-derived cells leads to a massive invasion of the forelimb bud by somite-derived YFP$^+$ cells (arrow).

FL, Forlimb; HL, Hindlimb; A, Anterior; P, Posterior; D, Dorsal; V, Ventral.

Scale bars: 500µm.
