## Supplemental Fig S2 for "Generation of a new Tbx6-inducible reporter mouse line to trace presomitic mesoderm derivatives throughout development and in adults"

**Supplemental Figure S2 (related to Figure 3)**

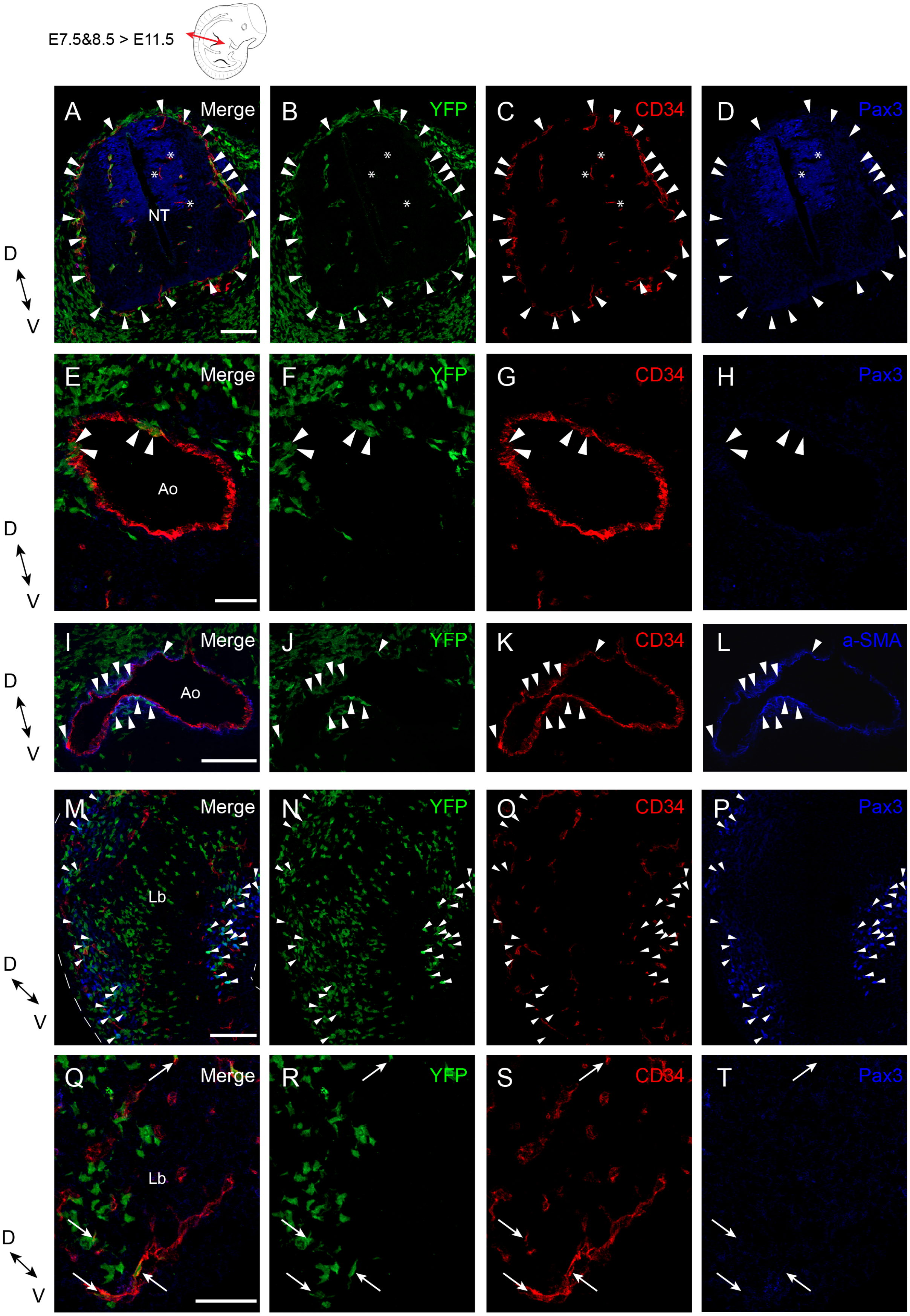

**Lineage tracing analysis of Tbx6-derived cells at E11.5 following tamoxifen induction at E7.5 and E8.5.**

(A-D) Close-up of the merge and single immunostaining pictures of the neural tube shown in ***Figure 3***. Merge (A), YFP (B), CD34 (C) and PAX3 (D) immunostaining pictures showing the contribution of the Tbx6-derived cells to the perineural vascular plexus (arrowheads). Of note, almost all the neural tube vascularization is derived from Tbx6(PSM-)-derived cells with few exceptions (asterisks). (E-H) Close-up of the merge and single immunostaining pictures of the dorsal aorta shown in ***Figure 3***. Merge (E), YFP (F), CD34 (G) and PAX3 (H) immunostaining pictures showing Tbx6-derived cells contributing to the endothelium of the dorsal aorta (arrowheads). (I-L) Close-up of the merge and single immunostaining pictures of the dorsal aorta obtained from an adjacent cryosection to the one shown in ***Figure 3,*** showing Tbx6-derived cells contributing to the smooth muscle around the aorta (arrowheads). Merge (I), YFP (J), CD34 (K) and α-SMA (L) immunostaining pictures. (M-P) Close-up of the merge and single immunostaining pictures of the forelimb bud shown in ***Figure 3***. Merge (M), YFP (N), CD34 (O) and PAX3 (P) immunostaining pictures showing the contribution of the Tbx6-derived cells to the myogenic progenitors colonizing the forelimb (arrowheads). (Q-T) Close-up of the merge and single immunostaining pictures of the forelimb bud tip shown in ***Figure 3***. Merge (Q), YFP (R), CD34 (S) and PAX3 (T) immunostaining pictures showing the contribution of the Tbx6-derived cells to the vascularization of the forelimb.

Ao, Aorta; Lb, Limb bud; NT, Neural Tube; D, Dorsal; V, Ventral.

Scale bars: 100μm in A-D, I-L, Q-T; 50μm in E-H, M-P
