## Supplemental Fig S3 for "Generation of a new Tbx6-inducible reporter mouse line to trace presomitic mesoderm derivatives throughout development and in adults"

### Supplemental Figure S3 (related to Figure 4)

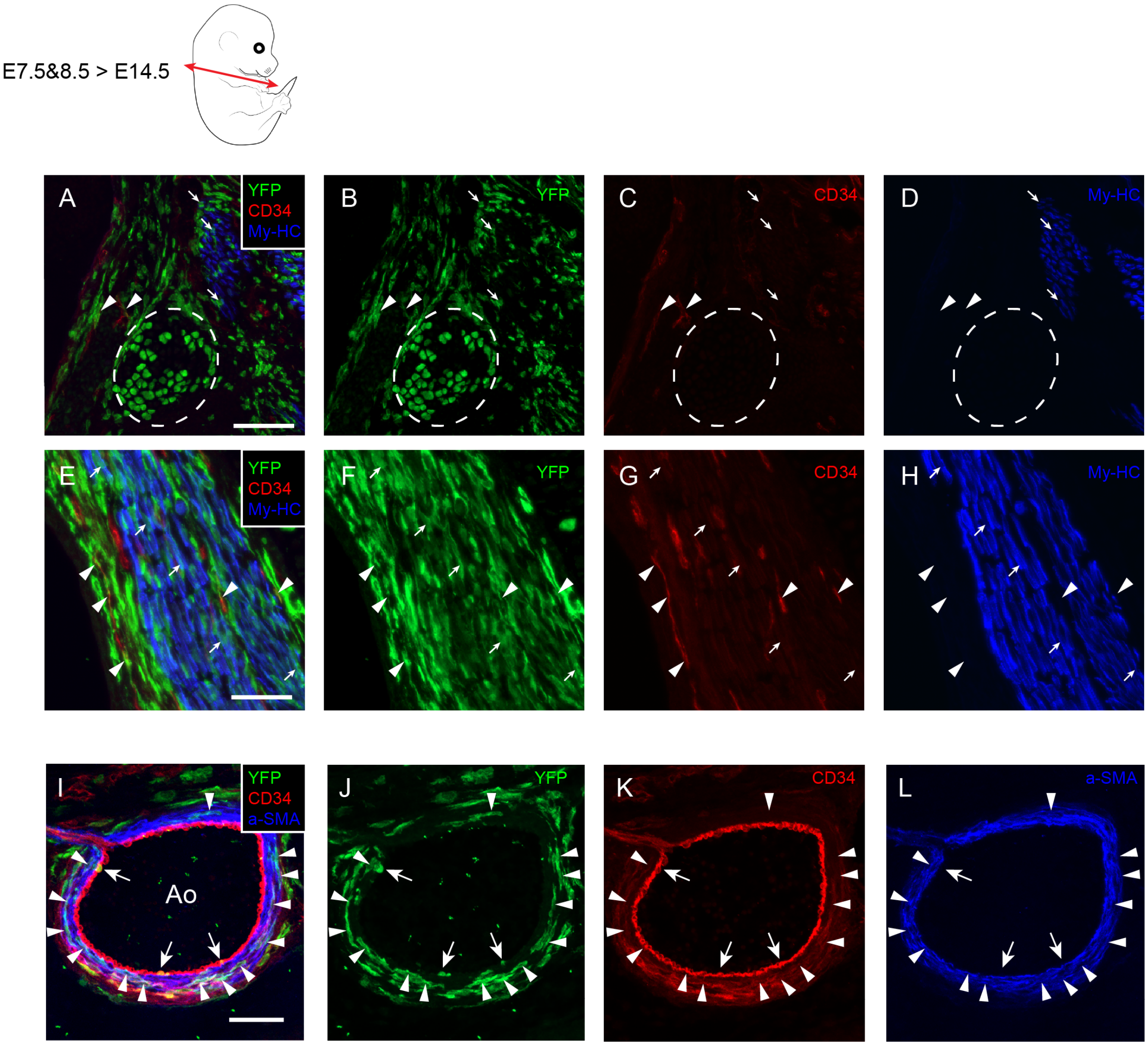


**Lineage tracing analysis of Tbx6-derived cells at E14.5 following tamoxifen induction at E7.5 and E8.5.**

(A-D) Close-up of the merge and single immunostaining pictures shown in ***Figure 4B***. Merge (A), YFP (B), CD34 (C) and MyHC (D) immunostaining pictures showing Tbx6-derived cells contributing to the vascularization (arrowheads) and muscles (arrows) of the trunk. Of note, some Tbx6-derived cells started to condense to form the future vertebrae (encircled area). (E-H) Close-up of the merge and single immunostaining pictures shown in ***Figure 4C***. Merge (E), YFP (F), CD34 (G) and MyHC (H) immunostaining pictures showing Tbx6-derived cells contributing to the vascularization (arrowheads) and muscles (arrows) of the limb. (I-L) Close-up of the merge and single immunostaining pictures of the dorsal aorta obtained from an adjacent cryosection to the one shown in ***Figure 4***, showing Tbx6-derived cells contributing to the smooth muscle around the aorta (arrowheads) and to a less extend to the endothelium of the aorta (arrows). Merge (I), YFP (J), CD34 (K) and α-SMA (L) immunostaining pictures.

Ao, Aorta.

Scale bars: 100µm in A-D; 50µm in E-L.
