## Supplemental Fig S4 for "Generation of a new Tbx6-inducible reporter mouse line to trace presomitic mesoderm derivatives throughout development and in adults"

**Supplemental Figure S4 (related to Figure 5)**

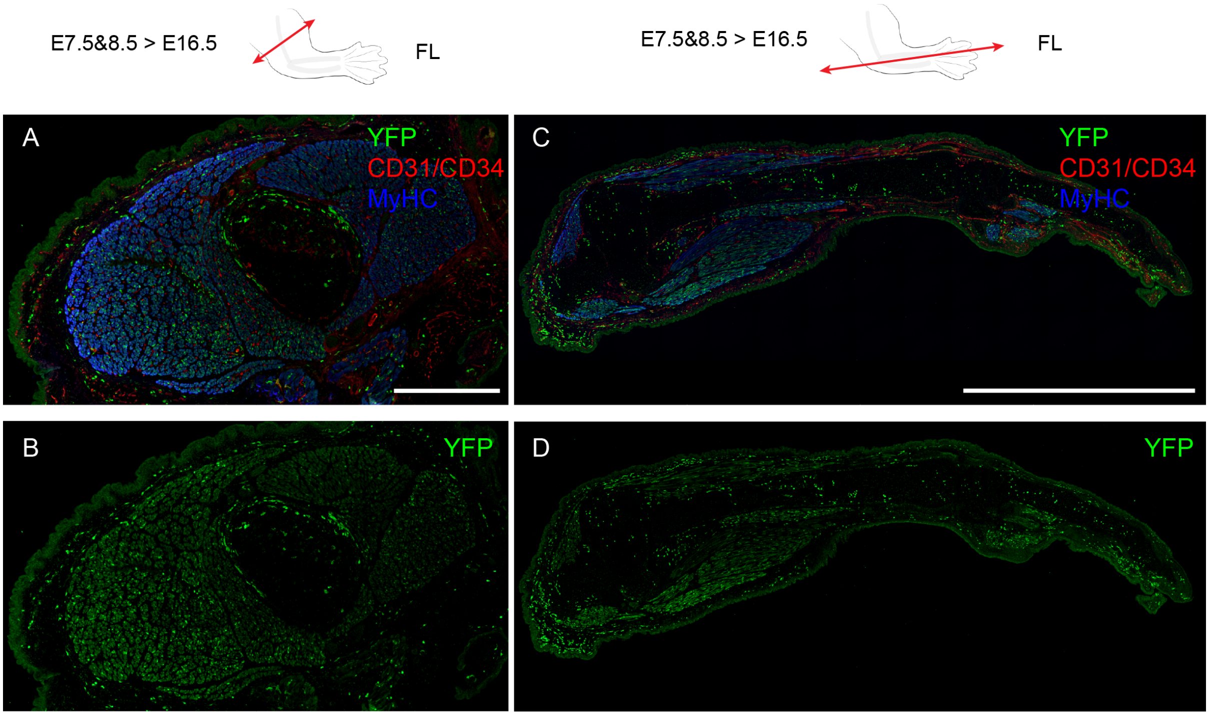


**Lineage tracing analysis shows the contribution of Tbx6-derived cells to the forelimb muscles at E16.5.**

(A-B) Transversal section of the forelimb. Merge (A) and YFP (B) immunostaining pictures showing the massive contribution of Tbx6-derived cells to the forelimb muscles. (C-D) Longitudinal section of the forelimb. Merge (C) and YFP (D) immunostaining pictures showing Tbx6-derived cells contributing to all muscles.

Scale bars: 500µm in A-B; 2mm in C-D.
