## Supplemental Fig S5 for "Generation of a new Tbx6-inducible reporter mouse line to trace presomitic mesoderm derivatives throughout development and in adults"

**Supplemental Figure S5 (related to Figure 6)**

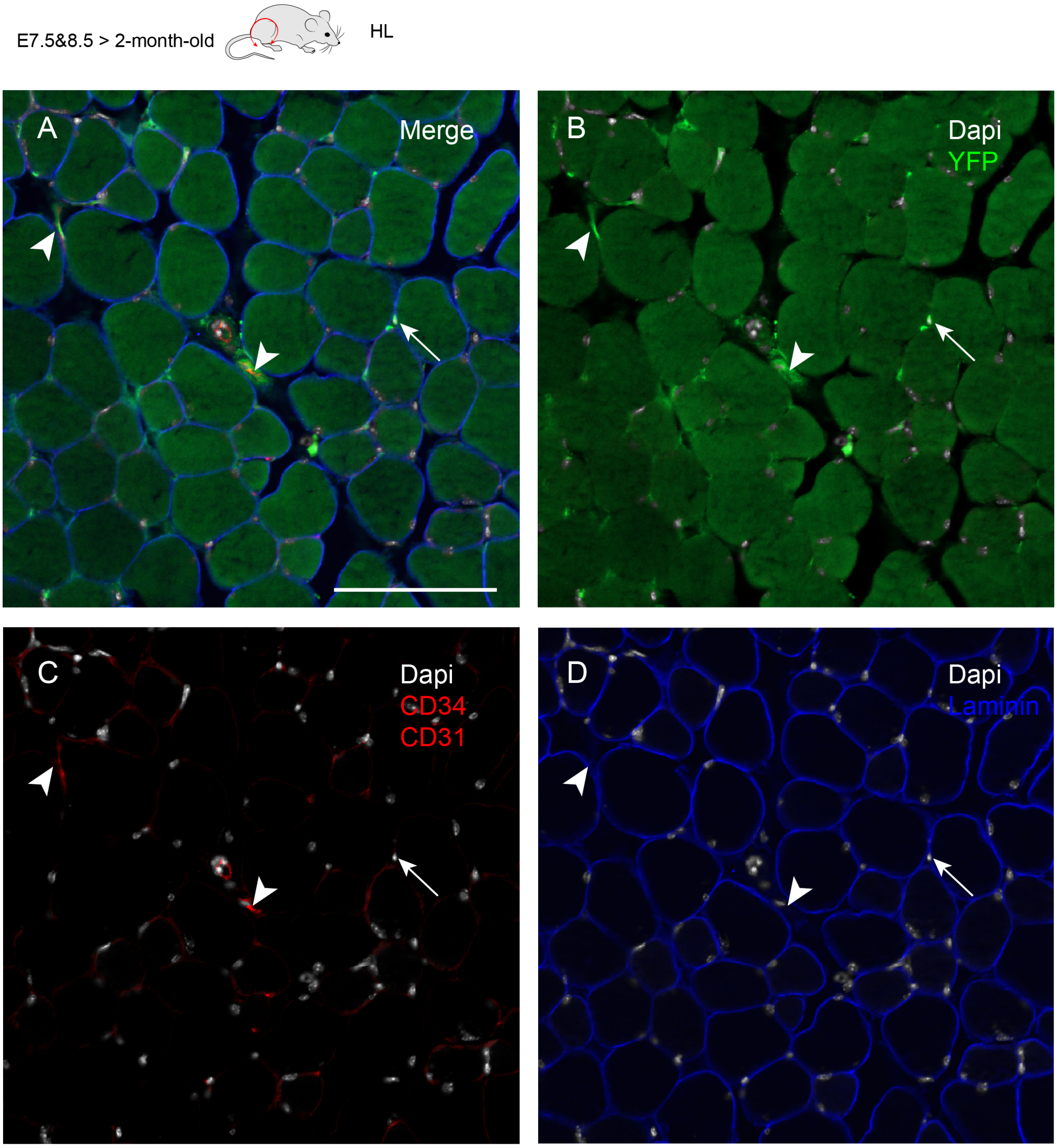


**Long-term lineage tracing shows the contribution of Tbx6-derived cells to the muscles and their vasculature in adults.**

(A-E) Transversal section of the hindlimb of a 2-month-old Tg(Tbx6_Cre/ERT2)/ROSA-eYFP mouse. Merge (A), YFP (B), CD34 and CD31 (C) and laminin (D) immunostaining pictures counterstained with DAPI showing Tbx6-derived cells contributing to the vascularization (arrowheads) and muscles (arrow).

Scale bar: 100µm.
