## Supplemental Table S1 for "Generation of a new Tbx6-inducible reporter mouse line to trace presomitic mesoderm derivatives throughout development and in adults"

| | Trunk | | | | | | Limbs | | | | | |
|---|---|---|---|---|---|---|---|---|---|---|---|---|
| | E11.5 | | | E14.5 | | | E11.5 | | | E14.5 | | |
| | YFP$^+$ pop | YFP$^+$ in ECs | ECs in YFP | YFP$^+$ pop | YFP$^+$ in ECs | ECs in YFP | YFP$^+$ pop | YFP$^+$ in ECs | ECs in YFP | YFP$^+$ pop | YFP$^+$ in ECs | ECs in YFP |
| Nb of embryos | 8 | 8 | 8 | 8 | 8 | 8 | 8 | 8 | 8 | 8 | 8 | 8 |
| Mean | 8.25 | 4,3 | 0,7 | 8,08 | 7,6 | 1,3 | 5,00 | 3,6 | 0,4 | 3,86 | 3,4 | 1,1 |
| SEM | 1.50 | 0,67 | 0,07 | 1,30 | 1,39 | 0,10 | 0,83 | 0,53 | 0,02 | 0,78 | 0,68 | 0,16 |

**Table S1.** Percentages of the total YFP cell population (YFP$^+$ pop), the YFP$^+$ cells within the endothelial population (YFP$^+$ in ECs) and the endothelial cells within the YFP$^+$ population (ECs in YFP) in the trunk or limbs, at E11.5 and E14.5.
